# Application of extracellular flux analysis for determining mitochondrial function in mammalian oocytes and early embryos

**DOI:** 10.1101/626333

**Authors:** Bethany Muller, Niamh Lewis, Tope Adeniyi, Henry J Leese, Daniel Brison, Roger G Sturmey

## Abstract

1.

**Background:** Mitochondria provide the major source of ATP for mammalian oocyte maturation and early embryo development. Oxygen Consumption Rate (OCR) is an established measure of mitochondrial function. OCR by mammalian oocytes and embryos has generally been restricted to overall uptake and detailed understanding of the components of OCR dedicated to specific molecular events remains lacking.

**Results:** Here, extracellular flux analysis (EFA) was applied to small groups of bovine, equine, mouse and human oocytes and bovine early embryos to measure OCR. Using EFA, we report the changes in mitochondrial activity during the processes of oocyte maturation, fertilization, and pre-implantation development to blastocyst stage in response to physiological demands in mammalian embryos. Crucially, we describe the real time partitioning of overall OCR to spare capacity, proton leak, non-mitochondrial and coupled respiration – showing that while there are alterations in activity over the course of development to respond to physiological demand, the overall efficiency is unchanged.

**Conclusion:** EFA is shown to be able to measure mitochondrial function in small groups of mammalian oocytes and embryos in a manner which is robust, rapid and easy to use. EFA is non-invasive and allows real-time determination of the impact of compounds on OCR, facilitating an assessment of the parameters of mitochondrial activity. This provides proof-of-concept for EFA as an accessible system with which to study oocyte and embryo metabolism.

## 2. Introduction

Re-emergence of the importance of metabolism in health and disease has stimulated investigations of mitochondrial function in the earliest stages of mammalian development. Mitochondria are abundant in the mammalian egg (Reynier et al. 2001; Collado-Fernandez et al. 2012), with mitochondrial DNA (mtDNA) copy number rising during oocyte maturation (Cotterill et al. 2013). By contrast, in the early embryo, mitochondria do not replicate until the blastocyst stage or later (Shoubridge 2000; Hendriks et al., 2018); thus, the pool of oocyte mitochondria at the point of ovulation support the entire pre-implantation period of development. In the oocyte and cleavage stage embryo, mitochondria exist in an immature though functional form (Motta et al. 2000; Sathananthan & Trounson, 2000). Acquisition of a typical mitochondrial morphology does not occur until blastocyst formation, corresponding to a period of increased energy demand (Fridhandler & Pincus 1957; Houghton et al. 1996; Thompson et al. 1996; Sturmey & Leese 2003; Lopes et al. 2007).

Mitochondrial oxidative phosphorylation, measured as the oxygen consumption rate (OCR) (Brand & Nicholls 2011), is the largest contributor to cellular ATP demand during preimplantation development (Brinster 1973; Leese 2012; Sturmey 2003; Van Blerkom, 2011). A number of techniques have been used to measure OCR in oocytes and embryos, including the Cartesian diver (Mills & Brinster 1967), microspectrophotometry (Herlitz & Hultborn 1974), pyrene ultramicrofluorescence (Houghton et al. 1996, Thompson et al. 1996), scanning electron microscopy (Shiku et al. 2001; Goto et al. 2018) and micro-respirometry (Lopes et al. 2005; Obeidat et al. 2018). Combined, these approaches have been instrumental in defining the overall metabolism of oocytes and early embryos, and have yielded remarkably consistent data. Importantly, OCR of mammalian embryos has been reported to correlate with reproductive physiological outcomes including oocyte maturation (Tejera et al. 2011), embryo morphology (Shiku et al. 2001; Lopes et al. 2007; Goto et al. 2018), implantation potential (Tejera et al. 2012), and pregnancy rate (Lopes et al. 2007). Generally, ‘mid-range’ or ‘Goldilocks-range’ OCR values are associated with higher oocyte/embryo quality and viability (Leese et al. 2016).

Despite research into embryo metabolism, a more comprehensive picture of OCR in oocytes and embryos in terms of the components of mitochondrial oxygen flux has remained elusive. A major limiting factor has been the lack of appropriate technology since current methods are time-consuming and technically demanding. This has restricted their applicability to Assisted Reproductive Technology (ART) clinics or as a tool for high-throughput screening or research. Recently, the emergence of extracellular flux analysis (EFA) technologies developed by Seahorse Bioscience (Agilent Technologies) has had a significant impact on the study of metabolism in a range of systems (Pelletier et al. 2014). EFA permits the real-time measurement of metabolism and the systematic evaluation of components of cellular oxygen consumption. We have applied this method the measurement of OCR by oocytes from a range of mammalian species, including the human. We have then explored the components of OCR on bovine embryos using EFA and discovered that mitochondrial function changes post fertilisation and between the cleavage and blastocyst stages.

## 3. Materials and methods

### 3.1 *In vitro* production (IVP) of bovine embryos

IVP was performed as described previously (Orsi et al., 2004). Unless stated otherwise, all media were prepared in the laboratory. Ovaries were collected from a local abattoir on the day of slaughter and transported in PBS warmed to 39°C within 3 hours of slaughter. Ovaries were washed twice in warmed PBS, and follicles were aspirated into warmed Hepes-buffered M199 containing heparin. Oocytes with compact cumulus were matured in groups of 50 for 18-22hrs in 5% CO_2_, 20% O_2_ in 500μl M199 containing 10% FBS supplemented with FSH, LH, EGF and FGF (BMM) at 39°C (Orsi et al., 2004). Fertilization was performed by co-incubating 0.5 x 10^6^ sperm/ml with groups of 50 oocytes in 500μl Fertilization-TALP (Fert-TALP) (Lu et al., 1987) for 24 hours. Presumptive zygotes were then denuded by vortexing, and transferred into 20μl drops of synthetic oviduct fluid (SOF) containing BSA and amino acids in groups of 20. These were incubated at 39°C in 5% O_2_, 5% CO_2_, 90% N_2_.

Oocytes or embryos were selected at the desired stage following standard IVP (described above), with the exception of GV-stage oocytes which were cultured overnight in cycloheximide (10µg/ml) to synchronize their stage. OCR was measured in groups of 6 CEOs following 24 hour culture in the presence of either meiosis-II (M-II) inhibitor cycloheximide, to prevent GVBD and resumption of meiosis, or of maturation-promoting hormones, to support progression to M-II stage. Expected proportion of oocytes at M-II stage after IVM was assessed by three independent nuclear staining experiments (0.1% Hoechst in 100% ethanol) following 18-22h IVM (Supplementary Figure S7). Presumptive zygotes were selected after 9 hours co-incubation with motile sperm. In order to assess approximate timing for appearance of PN following fertilization under our experimental conditions, independent staining experiments were performed. Oocytes were stained using 0.1% Hoechst between 4 and 12 hours following addition of sperm. We determined in independent experiments that 9 hours of co-incubation with motile sperm resulted in the highest concentration of PN-stage zygotes (Supplementary Figure S4). Embryos were selected out at the appropriate time (2-4 cell D2-D3, 8-16 cell D3-D4, blastocyst D6-D8) based on visual morphological staging.

Oocytes were analysed in BMM, PN-stage zygotes in HEPES-buffered Fert-TALP, and embryos in HEPES-buffered SOF – with all stages being analysed in groups of 6 in 180μl media. Equine COCs were analysed in groups of 3. Oocytes were denuded using 0.1% hyaluronidase and 1 minute vortexing, or stripped to corona using a holding pipette.

### 3.2 *In vitro* maturation of equine oocytes

Equine COCs were collected from abattoir-derived ovaries and held overnight as previously described (Choi et al., 2006; Hinrichs, 2005; Lewis et al., 2016). IVM was performed for 30 h in groups of three COCs with compact cumulus directly in a Seahorse XFp Bioanalyser plate in 180µl of maturation media (M199 with Earle’s salts, 10% FBS, 25µg/ml gentamicin with 5 mU/ml FSH (Hinrichs, 2005)). Only COCs with compact cumulus were used for this experiment.

### 3.3 Preparation of mouse oocytes

CD-1 (Charles River, Tranent, UK) mice were maintained, superovulated and housed overnight with stud males for mating as previously described by Ruane et al. (2017). Denuded fail to fertilize M-II oocytes (n=8) were obtained 36h post-ovulation and cultured briefly (2-4h) in 50µl KSOM medium (Millipore) containing 0.4% BSA at 37°C, 5% CO_2_ under ovoil (Vitrolife, Göteborg, Sweden) prior to assay in the XFP in a single group of 8.

### 3.4 Preparation of human oocytes

Appropriate consent to donate to research was obtained from patients undergoing Controlled Ovarian Stimulation (COS) for Intra-cytoplasmic Sperm Injection (ICSI) treatment in a National Health Service (NHS) IVF clinic as previously described (Gelbaya et al., 2006; Roberts et al., 2010). Ethics approval was obtain from the Manchester University Hospitals Research Ethics Committee and all donations were in accordance with the Human Fertilisation and Embryology Authority (HFEA) research licence R0026. All oocytes (n=5) were denuded MII failed to fertilize (OPN) fresh oocytes obtained from a single patient 18 - 20 h post ICSI. All oocytes were previously cultured in 50ul GTL medium under Ovoil (Vitrolife, Göteborg, Sweden) at 37^0^C, 5% O_2_ and 6% CO_2_ prior to assay in the XFP, 20 – 24 h post ICSI in a single group of 5 oocytes.

### 3.5 Application of Seahorse XFp to measure oxygen consumption in oocytes and embryos

Sensor-containing Seahorse fluxpaks (Agilent Technology) were incubated overnight at 37°C in a non-CO_2_ humidified incubator. The minimum time for incubation accepted was 8 hours and maximum 36 hours. The sensor-containing fluxpak was calibrated for approximately 15 minutes as per manufacturer guidance. Upon completion, the pre-warmed cell plate containing biological material was loaded into the machine. Oocytes and embryos were analysed using a specialised protocol involving a 12 minute equilibration period upon loading the cell plate, and alternating between a 3 minute measurement period and a 1 minute re-equilibration period. The measurement period involves the lowering of a sensor-containing probe, which creates an airtight 2.3 µl microenvironment in which change in pressure in mmHg is measured over time. This is followed by a 1 minute period in which the probe is lifted, and the 180μl well re-equilibrates. Plate specific ‘blank’ cell-free wells containing culture medium are used to account for environmental changes and flux of oxygen in the absence of cells, and as such oxygen consumption rate (OCR) is given as a function of these blank cell-free wells, with the flux in oxygen accounting for plasticware, diffusion of atmospheric O_2_, and finally specimen consumption (Gerencser et al. 2009). Output of Seahorse was given as oxygen consumption rate (OCR) in pmol/min/well.

To confirm that the assay had no impact on ongoing development, the blastocyst rates of bovine embryos which had undergone basal EFA on D2 were compared to those which were moved into HEPES SOF and kept in non-gassed incubation for the entirety of the Seahorse analysis (approximately 1 hour) as a control. Blastocyst rate was assessed daily between D6 and D8. Blastocyst rates observed in both groups were similar to expected rates given our IVP protocols (data not shown).

Both mouse and human embryos were analysed in the same manner – using 8 oocytes and 5 oocytes per well respectively. For equine COCs, the procedure was as described above with the following exception. As IVM took place directly in the cell plate (3 COCs per well), it was removed from the incubator and placed in the analyser at two different time points (4 and 28 hours after the start of IVM; IVM + 4hours, and IVM + 28 hours). After each set of readings was complete the plate was placed back in the incubator.

### 3.6 Use of mitochondrial inhibitors

Mitochondrial inhibitors were dissolved in 100% ethanol at 1000x the working stock. These were stored at −20°C for up to 3 months. Inhibitors were diluted in warmed analysis media (BMM, TALP or SOF, as dependant on stage of analysis) within 30 minutes of starting the assay.

Optimisation was carried out on GV-stage oocytes to establish appropriate concentrations of mitochondrial inhibitors indicative of mitochondrial parameters (Supplementary Figure S3). This involved serial injections of (1) oligomycin, (2) FCCP, and (3) a combination of antimycin A and rotenone. The Seahorse-XFp (Agilent) recommended concentration of 1µM oligomycin was used as a starting point, as it has been previously established to be appropriate for most cell types (Agilent Seahorse XF Cell Mito Stress Test Kit User Guide, 2017). This was tested in addition to 1.25, 2 and 3 μM. FCCP was tested at a variety of concentrations, using a titration of low (0.25 to 2µM) and high ranges (2.5 to 7.5µM). Antimycin A/Rotenone were tested at three concentrations: 1.5, 2.5 and 3.75. The lowest effective concentration was selected for each drug. Drug concentrations established for oocytes were applied to embryos. Oligo and A/R concentrations were effective on embryos, however FCCP was deemed too high, reducing OCR compared to basal, and was reduced to 3.75 µM (Supplementary Figure S5).

Inhibitors were loaded into the cell plate such that each injection represents 10% of the total well volume: 20µl, 22µl, 25µl and 27µl. As such, working concentrations were 10x the desired well concentration.

### 3.7 Data interpretation and statistical analysis

Wave software (Agilent Technologies) was used to determine oxygen consumption in pmol/min/well. This was normalised to number of oocytes/embryos per well. The third basal point, deemed most stable, was used as point of comparison for all data presented as proportion of basal. Point of highest response was used for all analysis for mitochondrial inhibitors. GraphPad Prism was used for all statistical analysis, using a significance level of p<0.05. Unpaired t-test was used where two groups were being compared. One-way ANOVA was used where basal OCR in more than two groups were being compared, while two-way ANOVA was used when drug response was considered between stages. Number of wells are used as technical replicates. Arc-sine transformation was used for the analysis of all proportional data. Number of wells in addition to number of oocytes/embryos used at each stage is described in Table 1. All graphs are presented as mean ± SEM.

**Table 1.**
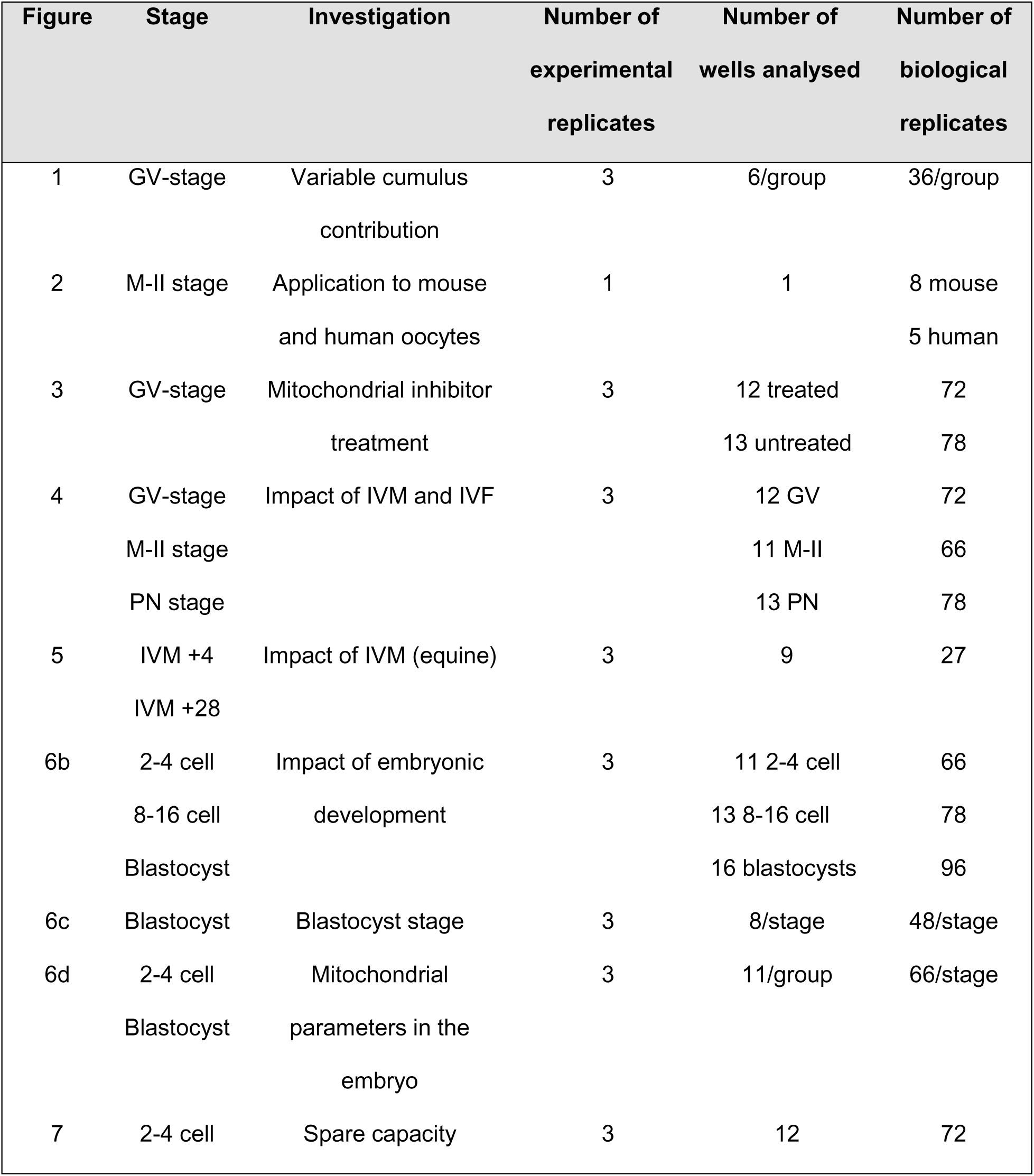
Experimental design. Experimental design for figures 1-5 is described, defining the variable of investigation, the developmental stage used and the number of samples per group. The ‘number of experimental replicates’ refers to pooled oocytes/embryos from the same ovary collection, ‘number of wells’ the number of Seahorse XFp wells analysed, and ‘number of biological replicates’ the number of oocytes/embryos used overall.

| Figure | Stage | Investigation | Number of<br>experimental<br>replicates | Number of<br>wells analysed | Number of<br>biological<br>replicates |
| --- | --- | --- | --- | --- | --- |
| 1 | GV-stage | Variable cumulus<br>contribution | 3 | 6/group | 36/group |
| 2 | M-II stage | Application to mouse<br>and human oocytes | 1 | 1 | 8 mouse<br>5 human |
| 3 | GV-stage | Mitochondrial inhibitor<br>treatment | 3 | 12 treated<br>13 untreated | 72<br>78 |
| 4 | GV-stage<br>M-II stage<br>PN stage | Impact of IVM and IVF | 3 | 12 GV<br>11 M-II<br>13 PN | 72<br>66<br>78 |
| 5 | IVM +4<br>IVM +28 | Impact of IVM (equine) | 3 | 9 | 27 |
| 6b | 2-4 cell<br>8-16 cell<br>Blastocyst | Impact of embryonic<br>development | 3 | 11 2-4 cell<br>13 8-16 cell<br>16 blastocysts | 66<br>78<br>96 |
| 6c | Blastocyst | Blastocyst stage | 3 | 8/stage | 48/stage |
| 6d | 2-4 cell<br>Blastocyst | Mitochondrial<br>parameters in the<br>embryo | 3 | 11/group | 66/stage |
| 7 | 2-4 cell | Spare capacity | 3 | 12 | 72 |

## 4. Results

### 4.1 Establishment of Extracellular Flux Analysis of bovine COCs

Initially, bovine oocytes at the germinal vesicle (GV) stage contained within intact cumulus complexes (COCs) were used to validate a method for the reproducible measurement of OCR. A linear relationship (r^2^=0.90; p=0.0042) was observed between the number of COCs and OCR, when comparing groups of between 1 and 25 COCs per well (Supplementary Figure S1). A comparison of group size (3 or 6) demonstrated that groups of 3 COCs were around the limit of detection of the assay system. Subsequent studies were therefore performed on groups of 6 COCs where possible.

### 4.2 Oxygen consumption in mammalian oocytes – the contribution of cumulus and the impact of *in vitro* maturation and *in vitro* fertilization

The contribution of cumulus cells to GV-stage oocyte respiratory activity was investigated by comparing the OCR of fully denuded oocytes (DO), corona-enclosed oocytes (CEOs) containing only the innermost 2-3 layers of cumulus cells, and fully intact COCs (Figure 1). OCR was significantly lower in DOs (0.44 ± 0.15 pmol/min/oocyte) and CEOs (1.68 ± 0.15 pmol/min/ooyte) compared to intact COCs (4.05 ± 0.75 pmol/min/oocyte). CEOs were selected for use in subsequent studies, to maintain the physiological state of oocytes, while keeping biological variation to a minimum.

**Figure 1.**
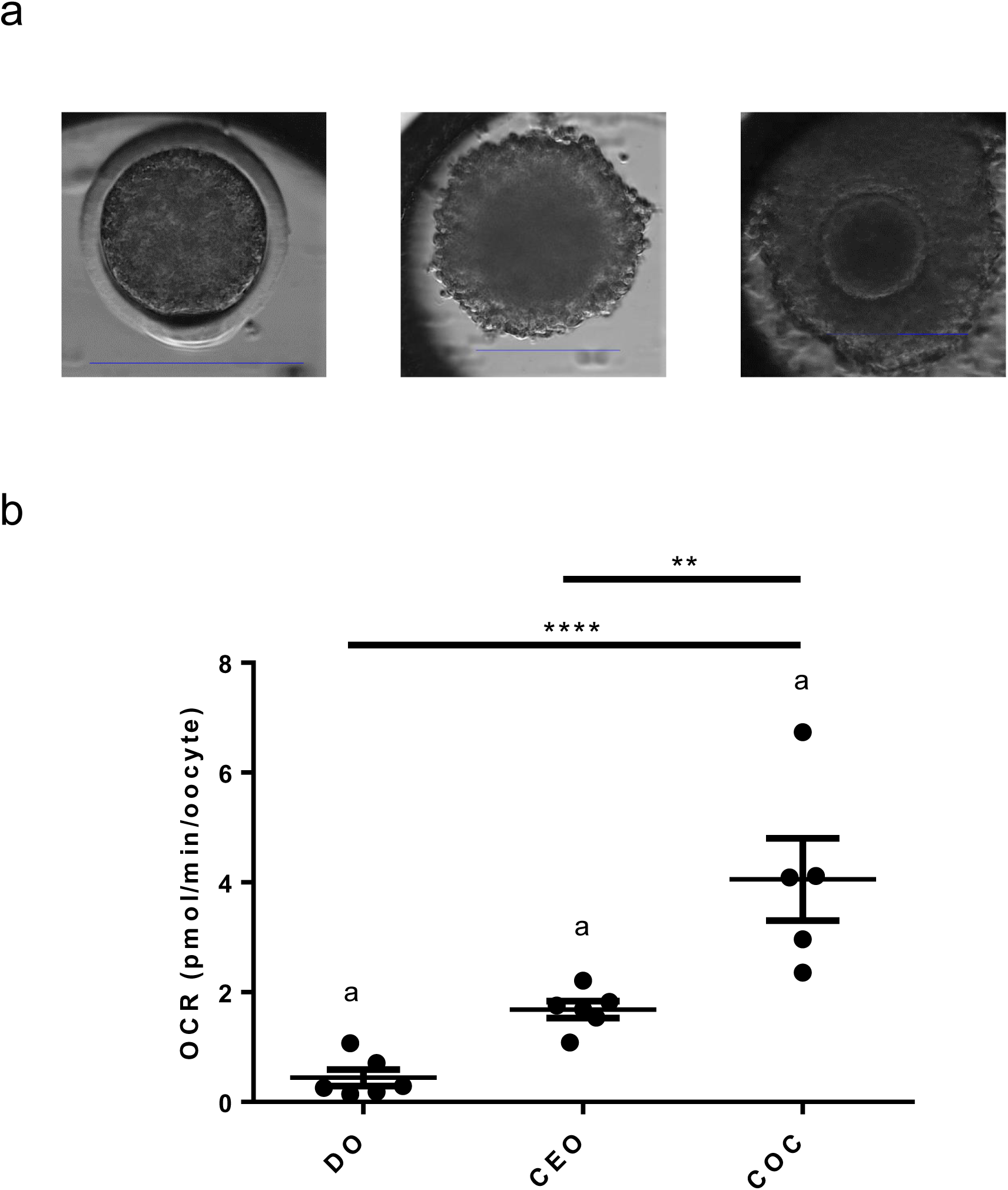
Contribution of the cumulus to bovine oocyte oxygen consumption at the GV-stage. (a) Time-lapse images of a fully denuded oocyte (DO), corona-enclosed oocyte (CEO) and intact cumulus oocyte complex (COC). Scale bars depict 150µm. (b) Basal OCR of DO, CEO and COCs. Data presented as mea n ± SEM, representing data from 6 wells (36 oocytes) per group. ** indicates p<0.01, **** indicates p<0.001. **a** indicates significant consumption of oxygen (p<0.05) using Wilcoxon Signed Rank Test.

EFA was then applied to small groups of mouse and human denuded M-II oocytes in order to demonstrate the applicability of the system across species. A high rate of basal OCR was observed in both species compared to bovine with 0.98 and 4.01 pmol/min/oocyte in mouse and human oocytes respectively (Figure 2). However, due to limited availability of material and the sensitivity limits of the system, the data represents a single well of oocytes thus demonstrating proof of principle but not allowing quantitative comparison.

**Figure 2.**
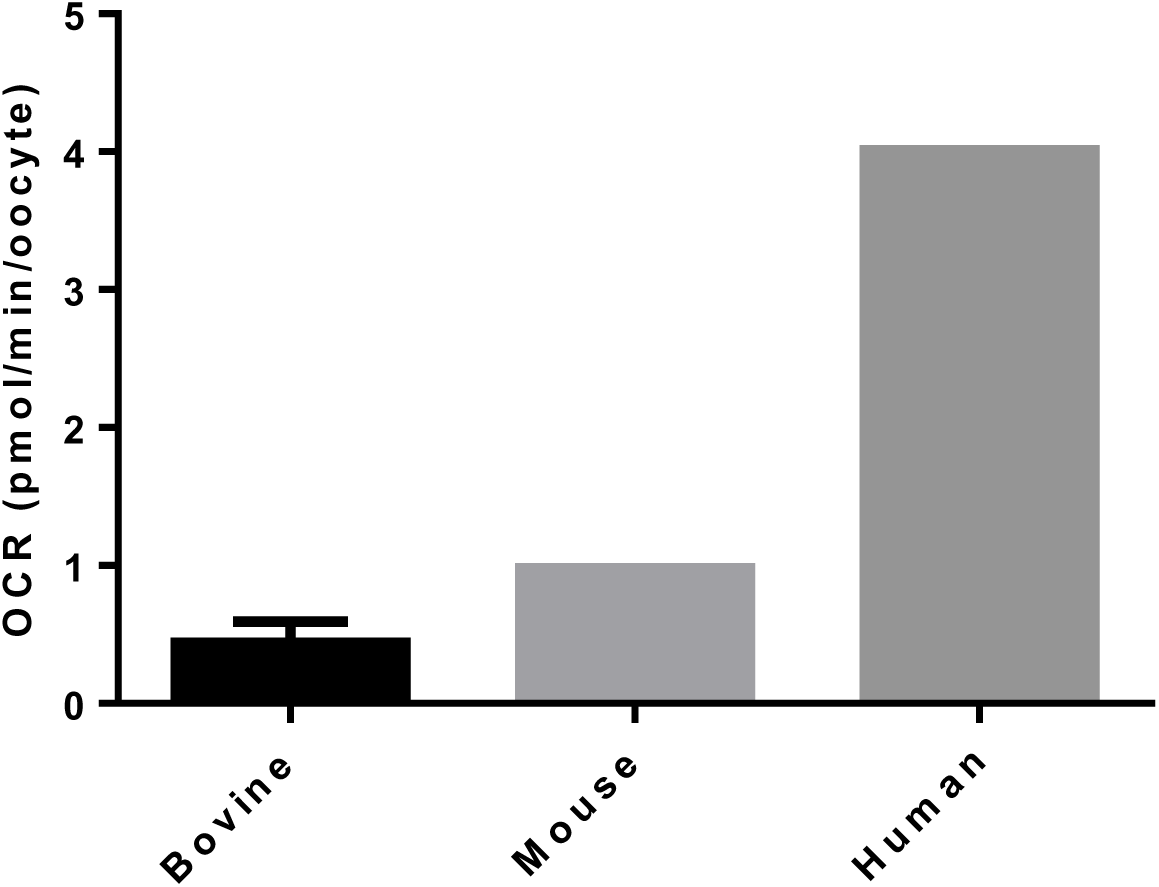
Basal OCR in denuded M-II mouse and human oocytes compared to GV-stage denuded bovine oocytes. Basal OCR of bovine (n=6, representative of 36 oocytes), mouse (n=1, representative of 8 oocytes) and human (n=1, representative of 6 oocytes) are indicated. Data is shown as mean ± SEM.

In order to measure the components of oxygen consumption of bovine CEOs, it was necessary to optimise the concentration of a series of mitochondrial inhibitors used to disrupt mitochondrial function (Supplementary Figure S3). Oligomycin inhibits ATP-synthase and can be used to indicate the proportion of O_2_ consumption directly coupled to ATP generation. Carbonyl cyanide-4-(trifluoromethoxy)phenylhydrazone (FCCP) is a mitochondrial uncoupler which dissipates the proton gradient between the inter membrane space (IMS) and the matrix allowing the measurement of maximal OCR. Added together, Antimycin A, a complex III inhibitor, and rotenone, a complex I inhibitor, combine to inhibit the ETC entirely; thus the proportion of OCR that is insensitive to the combined addition of Antimycin A and Rotenone (A/R) is considered to be non-mitochondrial in origin. 1µM oligomycin, 5µM FCCP and 2.5µM A/R gave the maximal response without killing the CEOs. The OCR fell to 43.9 ± 4.1% of basal OCR in response to inhibition of ATP synthase with oligomycin, rose significantly to 68.4 ± 21.0% above basal upon disruption of the proton gradient with FCCP, and fell to 22.7 ± 5.5% of basal OCR after the combined addition of A/R which totally blocks mitochondrial function (Figure 3). The per-well difference between non-mitochondrial and coupled OCR, proton leak, was 21.8 ± 6.2% (Figure 3c). These same concentrations were applied to mouse and human oocytes to mitochondrial inhibitors, and demonstrated that OCR was indicative of oxidative phosphorylation due to the fact that responses were observed to some extent despite lack of optimization of concentrations (Supplementary Figure S2).

**Figure 3.**
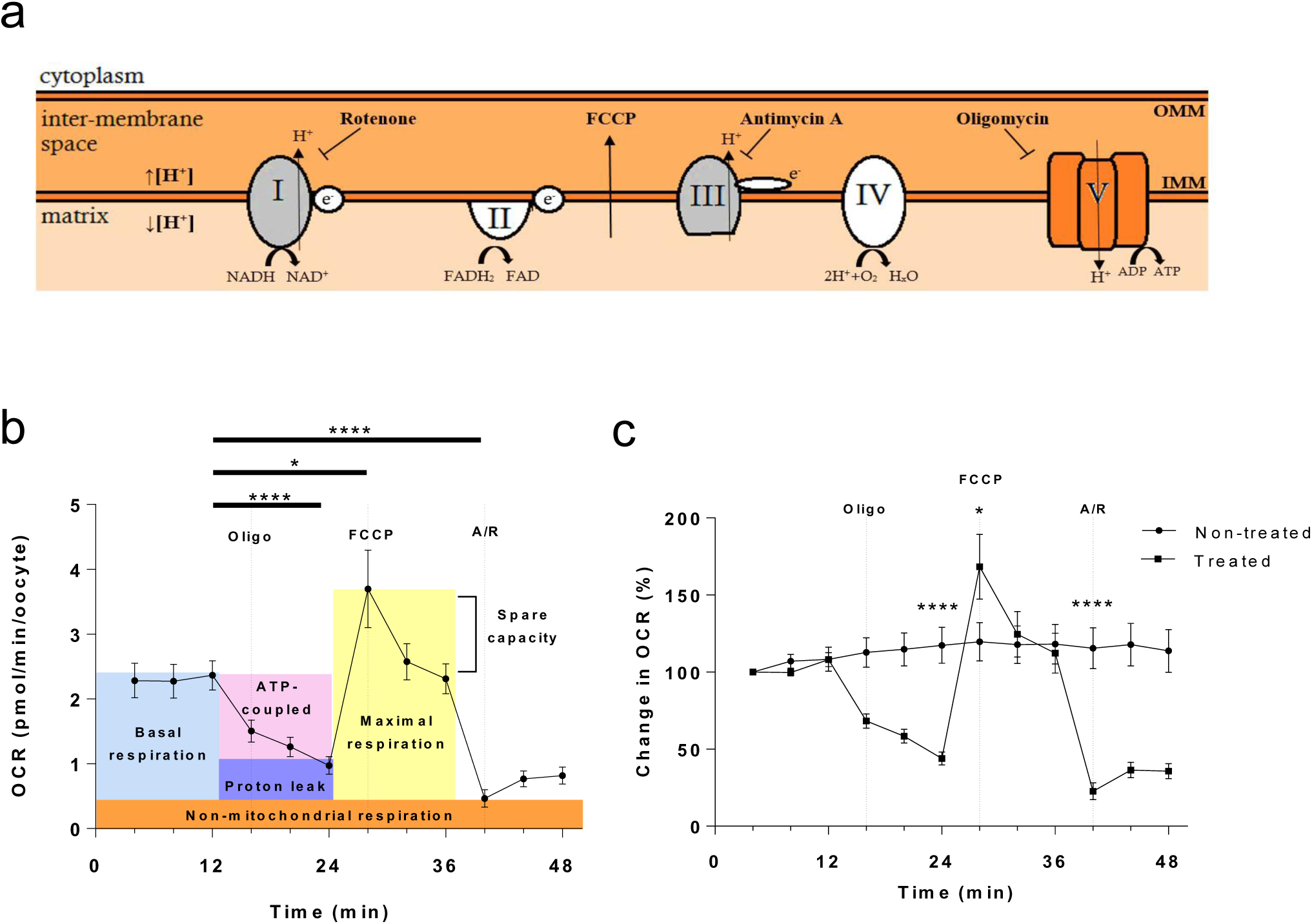
Application of mitochondrial inhibitors oligomycin, FCCP and antimycin A/rotenone to GV-stage oocytes. (a) Indicates the drug targets of Oligomycin, FCCP, Antimycin A and Rotenone. Oligomycin blocks ATP-synthase, FCCP dissipates the proton gradient between the matrix and inner membrane space, and antimycin A and rotenone inhibit complexes III and I respectively. (b, c) The response of GV-stage CEOs to the sequential addition of 1μM oligomycin, 5μM FCC and 2.5μM A/R, depicted as (b) OCR (pmol/min/oocyte) or (c) as compared to control non-treated oocytes, shown as percentage of the third basal measurement. (mean ± SEM, representative of 12 wells (72 CEOs) treated and 13 wells (78 CEOS) untreated). * represents p<0.05 and **** represents p<0.0001.

Having established the feasibility of measuring the components of OCR in CEOs, we next sought to discover whether the allocation of oxygen components changed during oocyte maturation and fertilisation. There was no difference in OCR (Figure 4a) or respiratory chain constituents (Figure 4b) between GV and MII oocytes. The optimal timing for pronuclear (PN) formation was determined as 9 hours post sperm addition (Supplementary Figure S4). Presumptive zygotes at the PN stage with intact corona demonstrated a slight, non-significant increase in OCR to 2.97 ± 0.45 compared to mature oocytes (p=0.17) and more notably a much wider data spread. Further, maximal OCR of PN zygotes was significantly higher than in mature oocytes (Figure 4b).

**Figure 4.**
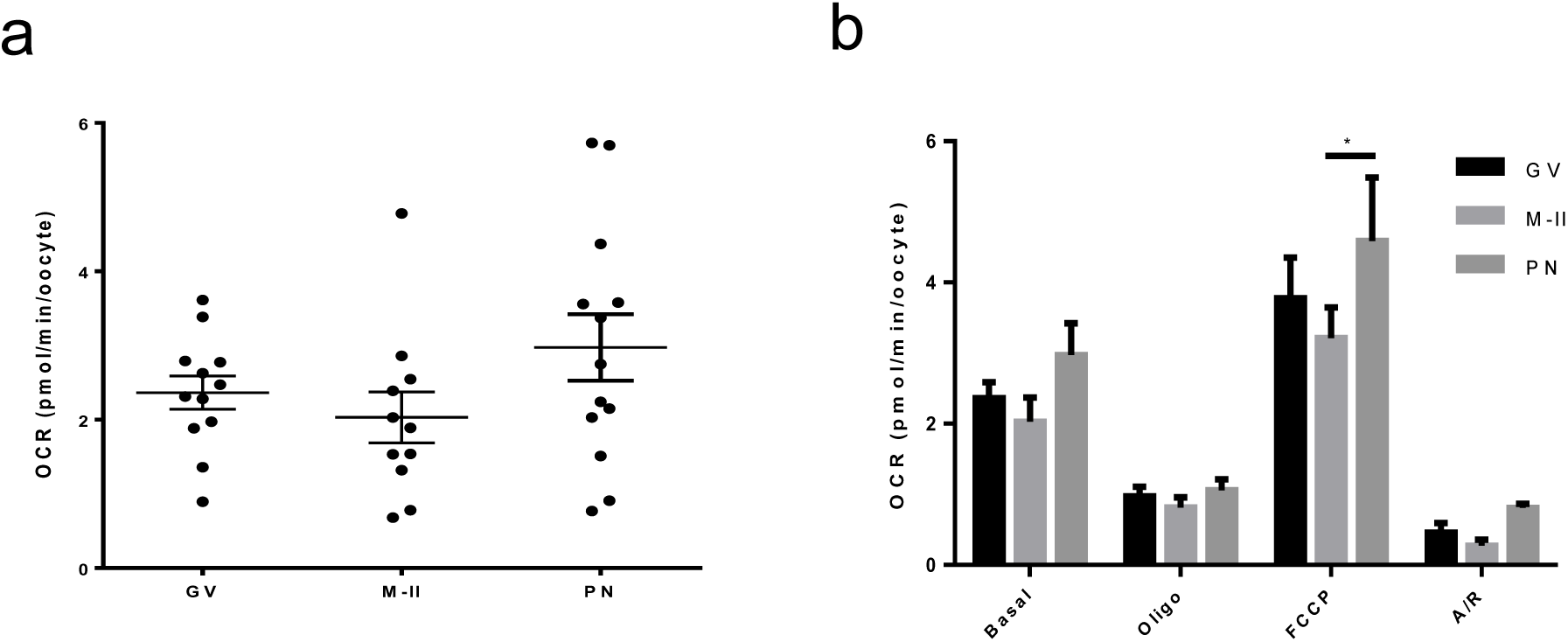
Oxygen consumption in bovine oocytes at GV, MII and PN stages. (a) Basal OCR of GV and MII-stage oocytes, and PN-stage zygotes (b) OCR of immature, mature and PN-stage zygotes in response to addition of 1μM oligomycin, 5μM FCCP and 2.5μM A/R (representative of 12 wells (72 oocytes), 11 wells (66 oocytes) mature and 13 wells (78 zygotes) respectively). All data are presented as mean ± SEM. All data presented as mean ± SEM, representative of 12, 11 and 13 wells of oocytes (72 and 66) and zygotes (78) at each respective stage. * depicts p<0.05.

In order to examine the relationship in another species, 27 equine COCs were analysed across three replicates at the beginning and end of a 30 hour maturation period. Basal OCR did not vary (Figure 5a) – with values of 38.1± 8.5 and 35.36 ± 9.7 pmol/COC/min after 4 and 28 hours respectively.

**Figure 5.**
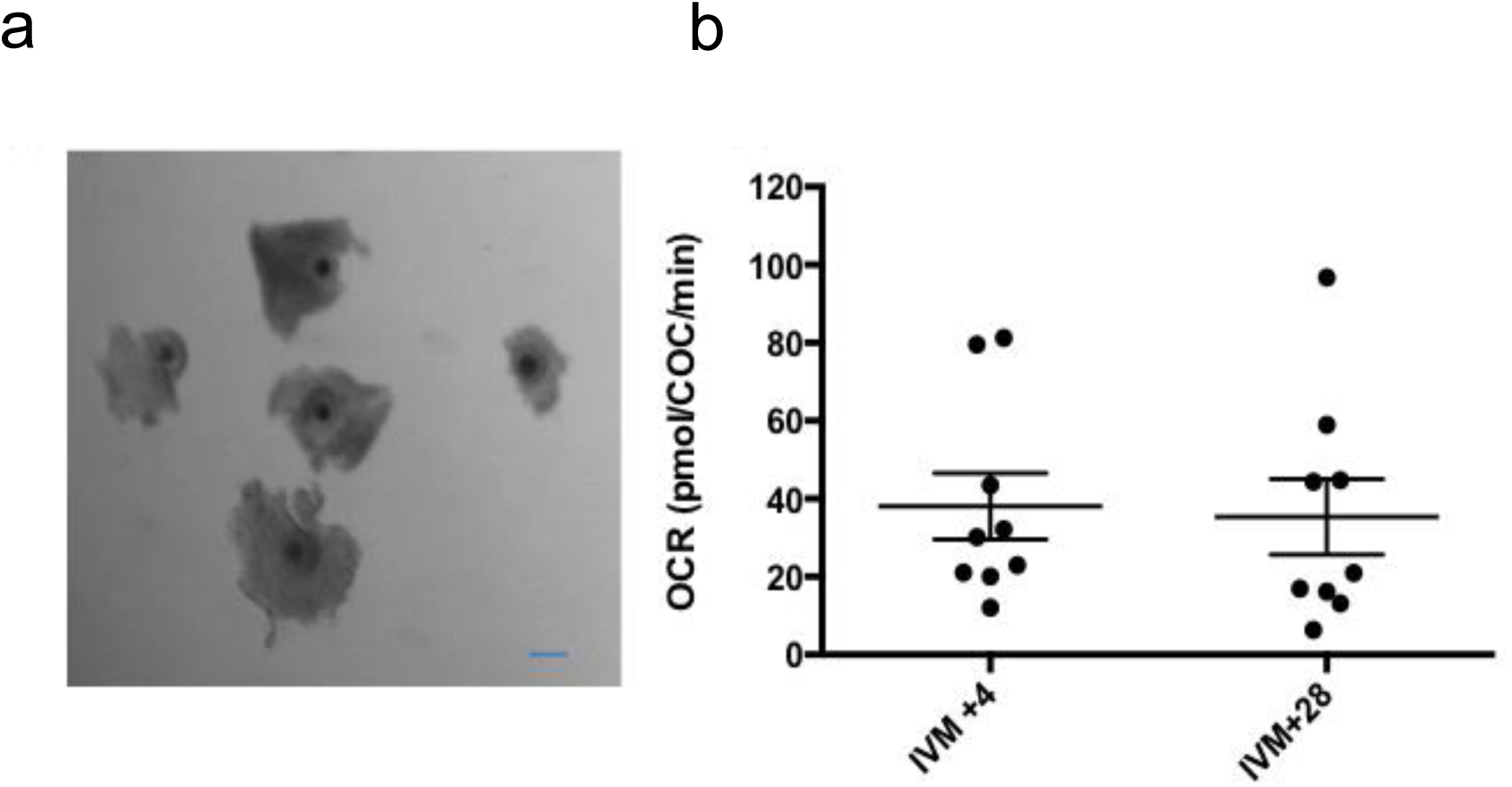
Basal OCR of equine cumulus enclosed oocytes at GV and MII stages of development. (a) Photomicrographs of equine COCs with compact cumulus. Scale bar depicts 100 µm (b) Basal OCR measured at 4 and 28 hours after the initiation of the 30 h in vitro maturation period (n=9 wells, representative of 27 COCs). Oocytes were cultured in Seahorse XFp plates between measurements. All data are presented as mean ± SEM.

### 4.3 Oxygen consumption in bovine embryos during preimplantation development *in vitro*

We next tested whether the Seahorse anlyser could record OCR by pre-implantation embryos. In the bovine, there was a gradual increase in OCR as embryos progressed through the cleavage stage - with OCRs of 0.54 ± 0.12 and 0.74 ± 0.07 pmol/min at the 2-4 cell and 8-16 cell stages respectively, followed by a significant increase at the blastocyst stage to 0.85 ± 0.08 pmol/min/embryo (p=0.02) (Figure 6b). When the OCR was compared between early and expanded blastocysts, there was a small increase which approached significance (p=0.066) (Figure 6c). Importantly, embryo viability was not altered by measurement of basal OCR since embryos that underwent 1 hour analysis were able to generate blastocysts with the same efficiency as equivalent controls (Supplementary Figure S5).

**Figure 6.**
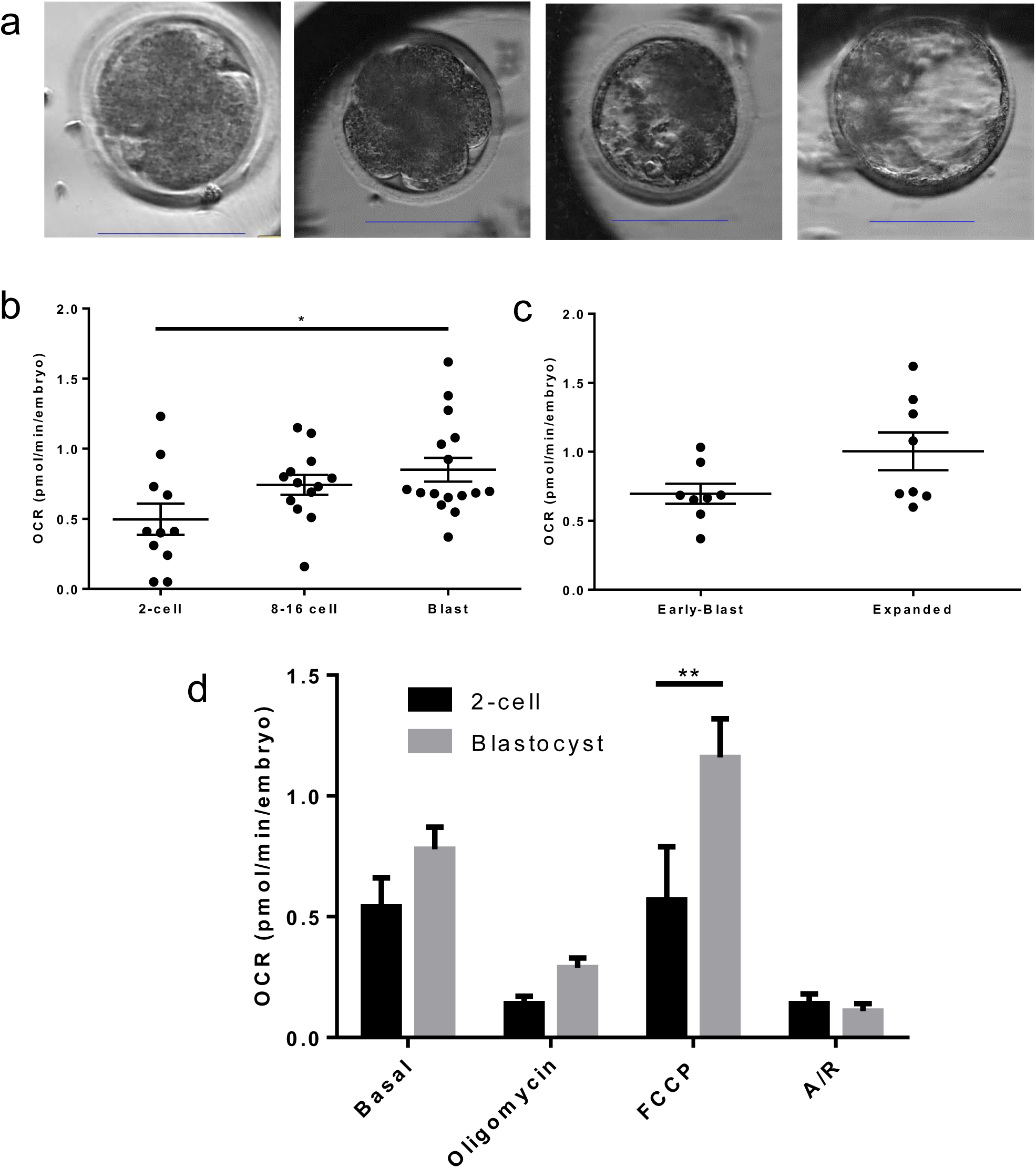
Oxygen consumption in embryos at cleavage and blastocyst stages. (a) Indicative time-lapse images of bovine embryos at 2-cell, 8-16 cell, full and expanded blastocyst stages. Scale bar depicts 100 µm. (b) Basal OCR in 2-4 cell, 8-16 cell and blastocyst stage embryos (representative of 66 2-4 cell embryos, 78 8-16 stage embryos, and 96 blastocysts). (c) OCR of early compared to expanded blastocysts (representative of 48 of each group). (d) OCR in response to inhibitors oligomycin, FCCP and A/R added sequentially at the 2-cell and blastocyst stage embryos (11 wells, 66 embryos per group). All data are presented as mean ±SEM. * indicates p<0.05, ** p<0.01.

Having measured basal OCR, the components of respiration were analysed in embryos. In initial experiments, the concentrations of mitochondrial inhibitors oligomycin and antimycin/rotenone used for oocytes were were found to be effective on embryos. By contrast, FCCP required further optimisation for use on embryos (Supplementary Figure S6). Using the revised concentrations of mitochondrial inhibitors, maximal respiration was significantly higher in blastocyst stage embryos comared to early cleavage (p=0.001), however OCR values in response to oligo and A/R indicative of coupled and non-mitochondrial respiration were unchanged between the stages (Figure 6d).

Whist examining the components of OCR by 2 cell embryos, we observed a sub population that did not respond to FCCP, indicative of a lack of respiratory spare capacity. This was an unexpected finding which was explored in more detail (Figure 7) which led to the finding that cleavage stage bovine embryos that failed to respond to FCCP had a significantly higher basal OCR (Figure 7a, b). When 36 2-4 cell embryos were allocated into groups depending on whether they had ‘low’ (0.22-0.43 pmol/min) or ‘high’ (0.64-0.81 pmol/min) OCR, the resulting blastocyst rates for those with ‘low’ OCR were slightly higher than those with ‘high’ (p=0.28) (Figure 7c).

**Figure 7.**
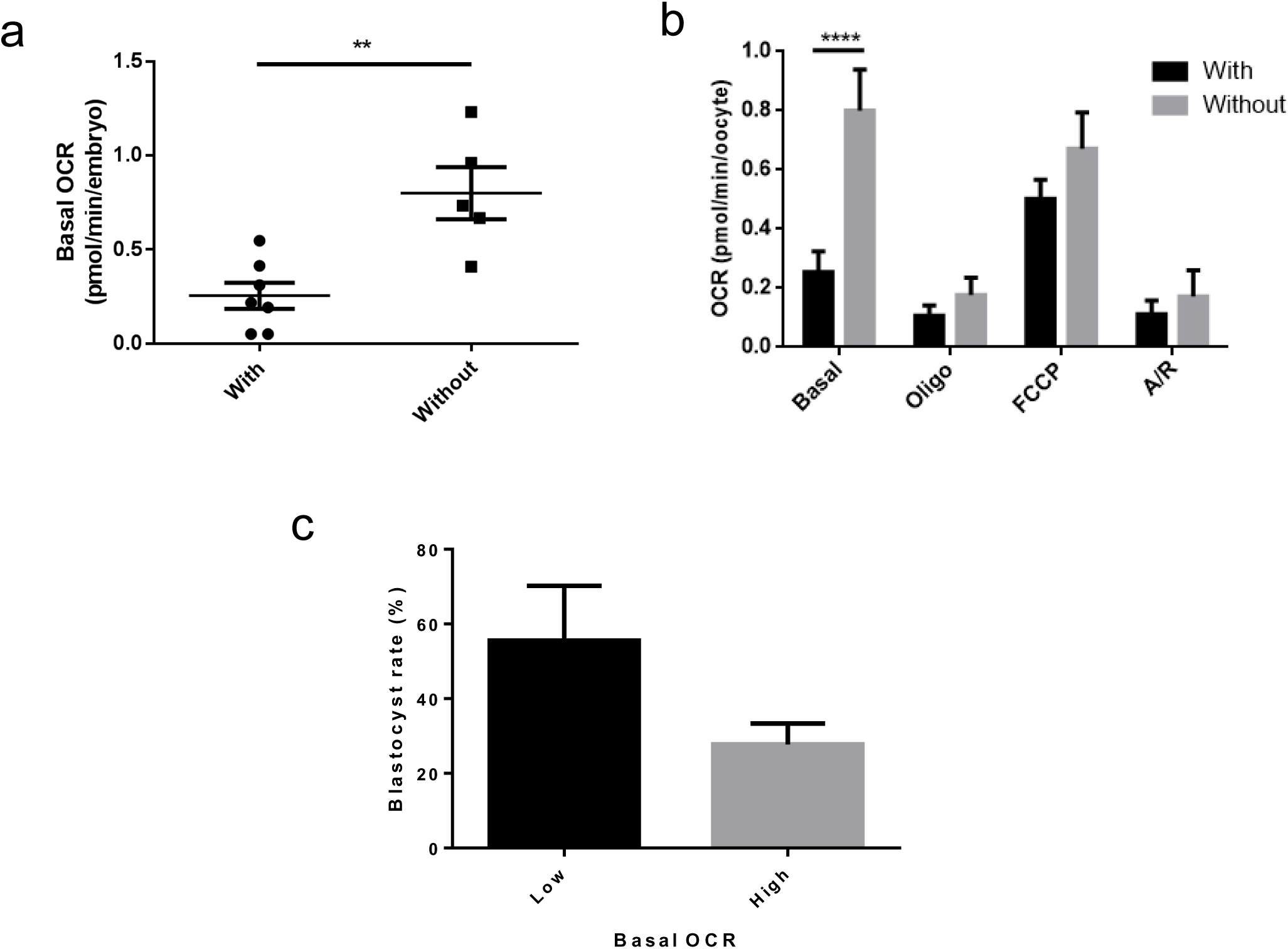
2-4 cell embryo mitochondrial activity. Early cleavage stage embryos showed variable response to FCCP. (a) Basal respiration (pmol/min/embryo) is shown for those with and without spare capacity (n=7 and n=5 respectively). (b) Drug response is shown across low and high groups. ** indicates p<0.01, **** indicates p<0.0001. (c) Blastocyst rate in embryos with low (ranging from 0.22-0.43 pmol/min/embryo) and high (ranging from 0.64-0.81 pmol/min/embryo) basal OCR at the cleavage stage (n=1, representing 36 blastocysts).

Bovine embryos, at all stages consumed significantly less oxygen than corona-enclosed oocytes and zygotes in terms of basal (Figure 8a) and maximal respiration (Figure 8b). However when the contributions of ATP-coupled, non-mitochondrial, proton leak and spare capacity to overall oxygen consumption across CEOs and embryos at key developmental stages were measured, there were no significant differences across development (Figure 8c).

**Figure 8.**
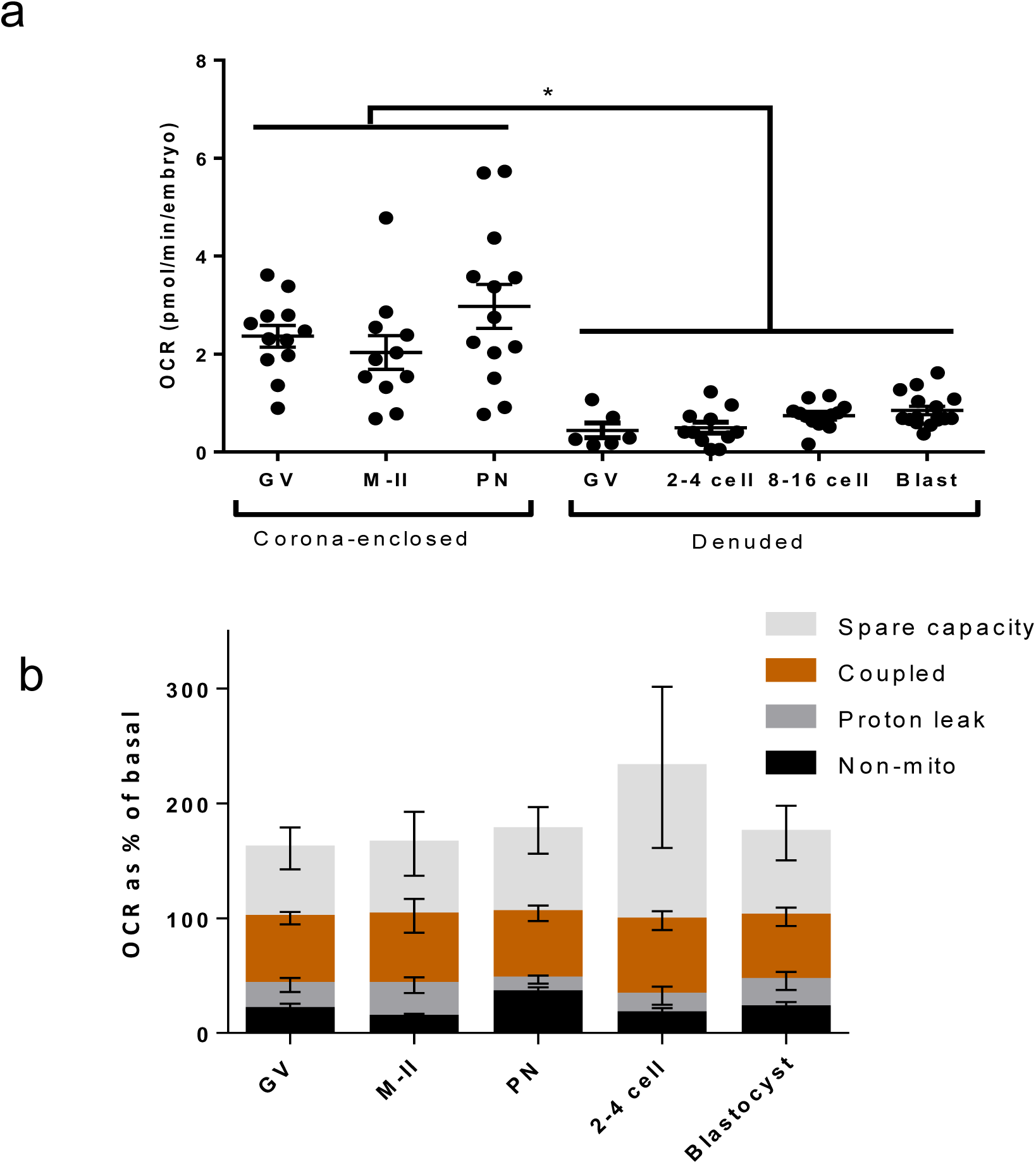
Summary of oxygen consumption across pre-implantation embryogenesis. (a) OCR of oocytes and embryos across pre-implantation development (representative of 72 GV-stage oocytes, 84 mII-stage oocytes, 78 PN-stage zygotes, 66 2-4 cell embryos, 78 8-16 stage embryos, and 96 blastocysts). (b) OCR in oocytes, PN-stage zygotes, 2-4 cell stage and blastocyst stage embryos in response to oligomycin, FCCP and A/R as indicative of mitochondrial parameters as a proportion of basal OCR (representative of 72 immature, 66 mature oocytes, 78 PN-stage zygotes and 66 embryos early cleavage and blastocyst stages). All data are presented as mean ± SEM. * indicates p<0.05.

## 5. Discussion

The metabolic processes supporting preimplantation development have been studied for more than 50 years (Brinster 1973; Leese 2012) with the aim of gaining fundamental understanding of the early mammalian embryo and identifying markers of embryo phenotype. While there is a mature set of data describing embryo metabolism, the assays used are technically complex and require dedicated equipment sensitive enough to measure very small quantities of biological material. In particular, the measurement of OCR, which is a marker of global oxidative metabolism, has only been reported for oocytes and embryos by a handful of laboratories. Here, we describe the first use of EFA as an accessible system which can measure oxygen consumption and its components in small groups of mammalian oocytes and early embryos.

### 5.1 Application of EFA to investigate the bovine cumulus-oocyte complex

After methods validation, the contribution of the cumulus cells to bovine oocyte respiration was determined. The mammalian oocyte is supported by close association with granulosa-derived somatic cumulus cells. The metabolic co-operation between cumulus and oocyte is vital (Eppig 1991) with cumulus cells being responsible for supplying key nutrients to the oocyte in the final phase of oocyte maturation (Sanchez-Lazo et al. 2014; Thompson et al. 2007; Leese & Barton 1984). Cumulus cells are preferentially glycolytic (Clark et al. 2006) and should therefore have only a minimal effect on oxygen consumption. However, we observed significantly lower OCR in denuded oocytes compared to COCs containing full cumulus contribution, indicating that the presence of cumulus cells does impact on the level of oxygen consumption in bovine oocytes. The data showing a significant difference between CEOs and COCs suggests that while each individual cumulus cell makes minimal contribution to OCR, the effect is amplified when the many cells work together, likely due to increased levels of signalling molecules as well as enhanced nutrient supply. Overall, our data support the notion that cumulus cells have a significant impact on oocyte oxidative metabolism.

### 5.2 Application of Extracellular Flux Analysis to mouse, human and equine oocytes

Using equine, mouse and human oocytes, the suitability of EFA to determine OCR in a variety of species was confirmed. The data were reassuringly close to those values obtained by previous methods; for example in the human, viable oocytes have been reported to consume 0.37 pmol/min (Tejera et al. 2011); in the mouse, 0.21 pmol/min (Harris et al. 2009); and equine oocytes 3 pmol/min (Obeidat et al. 2018); values close to our own. In the present work, mouse, human and equine oocytes showed higher basal respiration than bovine oocytes, however, the mouse and human oocytes used had failed to fertilise, suggesting they may not have been of optimal quality. Equine COC OCRs were approximately 10 times higher than bovine COCs. While the equine COCs were collected via follicular scraping rather than aspiration and likely had increased numbers of cumulus cells attached, the trend is in agreement with a previous equine study by Obeidat et al. (2018) in which denuded oocytes were used.

### 5.3 Application of Extracellular Flux Analysis to dissect mitochondrial function of bovine oocytes following maturation and fertilisation

The mitochondrial drugs oligomycin, FCCP and antimycin/rotenone were added systematically to dissect the components of OCR in mammalian oocytes. The data revealed that approximately 60% of OCR is coupled to ATP synthesis while a small but significant component of OCR (20%), is utilised for non-mitochondrial purposes. Bovine CEO at the GV stage has significant respiratory spare capacity, consistent with previous observations in bovine oocytes (Sugimura et al. 2012) and presumably required to satisfy dynamic energy demands as the oocyte undergoes maturation. These data confirm that the systematic interrogation of components of OCR in mammalian oocytes can be achieved through EFA.

Overall OCR did not change significantly over the course of culture in maturation media for bovine or equine oocytes. By contrast, a decrease in OCR post-IVM has previously been observed in denuded bovine oocytes at GV and M-II stages (Sugimura et al. 2012), which may reflect metabolic quiescence following meiotic progression and arrest at the M-II stage. However, in denuded pig and mouse oocytes, the opposite has been reported (Sturmey & Leese 2003, Harris et al. 2009), with M-II oocytes being more active than GV. This may reflect a species-specific difference, cumulus contribution, or be influenced by IVM protocols. It has previously been demonstrated that *in vitro* and *in vivo* matured oocytes show differential OCR profiles (Sugimura et al. 2012).

In the corona-enclosed early zygote, there was a trend in the direction of increased basal OCR, and FCCP application led to a significant increase in maximal OCR compared to oocytes post-IVM. This pattern suggests an increase in mitochondrial activity and reserve capacity coincident with fertilisation. A peak in mitochondrial activity at fertilisation has previously been shown, first in the sea urchin (Heinecke & Shapiro 1989) and later in the bovine model (Lopes et al. 2010). It has been proposed that this phenomenon occurs in response to calcium oscillations (Campbell & Swann 2006) which are triggered by PLCζ at the time of fertilisation, and which promote oxygen consumption in oocytes (Dumollard 2003; Van Blerkom 2003). This rise in mitochondrial activity may be due to the ATP demands of chromosome reorganisation for PN formation and extrusion of the second polar body. Demonstration of the presence of a reserve capacity (the difference between maximal and basal OCR) is of note since mitochondrial biogenesis, the normal physiological response to increased ATP demand, does not occur at this stage. However, paternally-derived mitochondria will still be present since they are not degraded until early cleavage (Sutovsky 2014) and may contribute to the increased maximal capacity we observed. It is notable that following both IVM and IVF not all oocytes were at the expected developmental stage (Supplementary Figure S4 & S7). Since only 60-70% were expected to be fertilised, this strengthens our observation of an increase in mitochondrial activity at PN-stage.

### 5.4 Application of Extracellular Flux Analysis to investigate bovine embryo physiology

The OCR did not differ throughout the cleavage stages of bovine preimplantation embryo development. Such a pattern has been previously described for mouse (Houghton et al. 1996), cow (Thompson et al. 1996; Lopes et al. 2007), and pig (Sturmey and Leese 2003). Upon reaching the blastocyst stage, there was a significant rise in OCR (Figure 6) - in agreement with earlier reports (Fridhandler & Pincus 1957; Houghton et al. 1996; Thompson et al. 1996; Sturmey & Leese 2003; Lopes et al. 2007).

The components of OCR were measured in 2-cell embryos and blastocysts. Not unexpectedly, blastocysts exhibited a higher maximal respiratory rate compared to 2-cell cleavage-stage embryos. This increase in both basal and maximal OCR in blastocysts supports the notion of metabolic plasticity in the preimplantation embryo and provides the facility by which blastocysts can rapidly increase ATP synthesis. ATP demand increases at this stage to meet the needs for Na^+^/K^+^ ATPase activity, necessary for production and maintenance of the blastocoel cavity, to support a rise in protein synthesis, and allow for cellular differentiation (Biggers et al. 1988). Replication of mtDNA has been shown to be initiated at the blastocyst stage in the horse (Hendriks et al. 2018).

At the 2-4 cell stage of development, a high degree of heterogeneity in basal OCR and FCCP response was apparent and curiously, a significant proportion of cleavage stage embryos failed to respond to FCCP. In terms of the basal respiration, those cleavage-stage embryos which did not respond to FCCP had a significantly higher basal respiration. Thus, at the 2-cell stage, it was apparent that embryos fell into two distinct groups – one with low OCR and a spare capacity, and one with high OCR and no spare capacity. Nevertheless, the proportion of OCR ascribed to ATP synthesis and non-mitochondrial functions did not differ between the two groups. The reasons behind there being two distinct groups are not immediately clear. However, it has been proposed that the most viable embryos will be metabolically mid-range when distributions of individual values are plotted, while those that exhibit higher or lower metabolic activity are more likely to be stressed (Guerif et al. 2013; Leese et al. 2016). Arrest at the 2-4 cell stage is a common phenomenon, representing around embryos 10-15% in human ART (Betts & Madan 2008) and 15% in the bovine model (Matwee et al. 2000). A number of factors have been implicated in this developmental arrest including its coinciding with the pressures of embryonic genome activation (Artley et al., 1992), chromosomal abnormalities (Almeida et al., 1998), abnormal cleavage events (Burruel et al. 2014) and reactive oxygen species (ROS) generation and oxidative stress (Kimura et al. 2010; Favetta et al. 2007). A preliminary investigation into whether these different groups reflected developmental capacity showed that those in the ‘low’ OCR group were more likely to go on to form blastocysts, suggesting a physiological difference between the groups rather than an observation of timing of cleavage divisions. However, this experiment used ovaries collected on a single day and further work is required to elucidate this intriguing relationship and the mechanism(s) behind it. The use of 6 embryos grouped at random represents a limitation to this finding. The distinct groups observed implies either that the proportion of ‘low’ and ‘high’ OCR embryos affected the resultant reading, or that there might be a paracrine effect from particularly stressed over-active embryos. Our observation could also stem from the stochastic nature of the measurement – it has been demonstrated that each cytokinetic event of cleavage is associated with small peaks in mitochondrial activity (Tejera et al., 2016).

It is notable that the present values of basal OCR (Figure 8a) are within the range of what has been observed previously, though direct comparisons are difficult due to the differences in technical approach and reporting data. Denuded bovine 1-cell zygotes have been reported to consume approximately 1.78 pmol/min (Thompson et al. 1996), an increase compared to un-fertilized oocytes. Sugimura et al. (2012) reported figures of 0.26 and 0.18 pmol/min prior to and following IVM in denuded bovine oocytes while Lopes et al. (2010) found a value of 2.83 pmol/min in mature bovine oocytes and similar values in D3 embryos, with blastocysts undergoing a two to three-fold increase in respiratory activity, though this was affected by sex - more pronounced in female/male than female/male embryos, and by morphological quality (Lopes et al. 2005). Obeidat et al. (2018) recently reported denuded bovine oocyte respiration to be approximately 1.2 pmol/min and embryo to be 4.2 pmol/min, though stages were not clearly indicated. These previous studies have therefore reached similar conclusions to our own both in terms of raw numbers but also physiological trends such as a modest increase following fertilisation, and a major increase between cleavage and blastocyst stages.

### 5.5 Markers of mitochondrial activity across pre-implantation development

Basal and maximal OCR of embryos were significantly below that of cumulus enclosed oocytes at all stages analysed. This most likely reflects the interaction with cumulus that occurs between these stages, since the OCR of denuded oocytes was similar to that measured in early cleavage-stage (2-4 cell) embryos. The contributions of non-mitochondrial OCR, proton leak and coupled respiration expressed as proportions did not differ at any stage of preimplantation development indicating that while overall oxygen consumption might be changing, the components of respiratory function remained similar. We consider this a key finding since it indicates that while there are alterations in respiratory activity over the course of development in response to physiological demand, the overall efficiency of the process is unchanged; rather, that a change in mitochondrial number/mass, substrate availability or rate of activity of the ETC is occurring. Around the blastocyst stage, mitochondrial biogenesis begins (Hendriks et al. 2018), consistent with the increase in both basal and maximal OCR observed here. Further, reliance on different fuels (carbohydrates, fats or proteins) can alter mitochondrial activity by affecting the respiratory quotient, a term which describes the ratio of CO_2_ production to O_2_ consumption (Muoio 2016) – thus OCR may be a useful means for non-invasive mapping of changes in nutrient metabolism known to occur across embryogenesis (Leese 2012).

Throughout oocyte maturation and preimplantation embryo development, approximately 20% of OCR could be accounted for by proton leak, while about 60% was coupled to ATP formation. Although some minor variations in proportions were observed at the different stages, proton leak and the coupling to ATP production remained relatively constant. The figure of 20% for proton leak has been observed across a range of cell types (Brand 2000). It decreases superoxide production due to its effect on the proton gradient, and is therefore involved in the regulation of ROS production. The process of uncoupling thus has a protective role against ROS, observed for example when under oxidative stress (Cadenas 2018) and in ageing (Brand 2000). Coupling efficiency can vary significantly between different cells types, from as low as 30% to up to 90% (Brand & Nicholls 2011). This can vary as dependent on ATP demand or in order to regulate ROS levels (Cadenas 2018), and is thought to stem at least in part due to tissue and cell-type specific differences in mitochondrial structure (Woods, 2017).

Our observation of around 20% of basal OCR derived from non-mitochondrial processes was of particular interest. This figure is higher than for most somatic cells, which tend to give a figure around 10% (Brand & Nicholls 2011), although it was lower than previously determined in rabbit, mouse and bovine oocytes and embryos (Manes & Lai 1995; Trimarchi et al. 2000; Sugimura et al. 2012) – who reported values of approximately 25%, 23-30%, and 35-40% respectively. This is likely due to biological differences as well as experimental approach. Sources of non-mitochondrial oxygen consumption include cell-surface oxygen consumption from electron transport at the membrane and enzymatic ROS production, for example NADPH oxidase in the rabbit blastocyst (Manes & Lai 1995; Herst & Berridge 2007; Starkov 2008). This may relate to the involvement of ROS in redox regulation which may play essential physiological roles in, for example, oocyte activation, embryonic genome activation and hatching (Harvey et al. 2002). Cell-surface oxygen consumption has been reported to support rapidly proliferating tumour cells highly active in glycolysis (Herst & Berridge 2007), and might play a similar role in dividing embryos which exhibit aerobic glycolysis (Krisher & Pather, 2012).

Spare respiratory capacity was the function with the greatest variation. Overall, a large spare capacity was observed at the stages assessed, though the exception to this generalisation was the sub-population of embryos at the 2-cell stage that did not exhibit spare capacity. The spare capacity may be regulated by the contribution of cumulus cell mitochondria, changes in mitochondrial mass after fertilisation and at the blastocyst stage (Sutovsky 2014; Hendriks et al. 2018) and response to physiological demand. This could in part facilitate the peak of increased mitochondrial activity that occurs for example at fertilisation and with cleavage divisions (Tejera et al. 2016). Reserve capacity has also been demonstrated to be linked to increased cellular survival in cardiomyocytes and in fibroblasts (Hill et al. 2009; Nickens et al. 2013), ensuring cells have the capacity to provide more ATP should conditions require it.

Reserve capacity is facilitated by regulation of the TCA cycle and of complex II of the ETC, which respond to substrate availability (Pfleger et al. 2015). Metabolic sensors are used to set off this regulation, thus allowing the cell to meet demands and react to stressed conditions. We speculate that the high levels of spare capacity observed throughout may contribute to the metabolic and developmental plasticity that oocytes and embryos show during the periconceptual stage of development that is highly responsive to environmental conditions (Eckert & Fleming 2011). Environmental challenges to the embryo may present in the form of maternal diet or *in vitro* culture conditions – conditions under which metabolic activity has been shown to be altered (Sugimura et al. 2012; Leary et al. 2015).

### 5.6 Prospects for the future application of extracellular flux analysis

Overall, this research has shown that EFA can readily be used as a tool to measure oxygen consumption in real-time as a proxy for the function of mitochondria and their components (spare respiratory capacity, proton leak, non-mitochondrial and coupled respiration) in small groups of mammalian oocytes and preimplantation embryos. The approach is robust and rapid, and its automated nature limits room for operator-induced error and potential harm to the gametes and embryos. Crucially, analyses are non-invasive and do not impact on ongoing *in vitro* development. Future work may lead to optimisation of the Seahorse XFp system to enable higher sensitivity, allowing it to be applied to single oocytes and embryos, with the potential to screen individual embryos prior to transfer in clinical and farm animal IVF.

## Supplementary Figures

**Figure S1.**
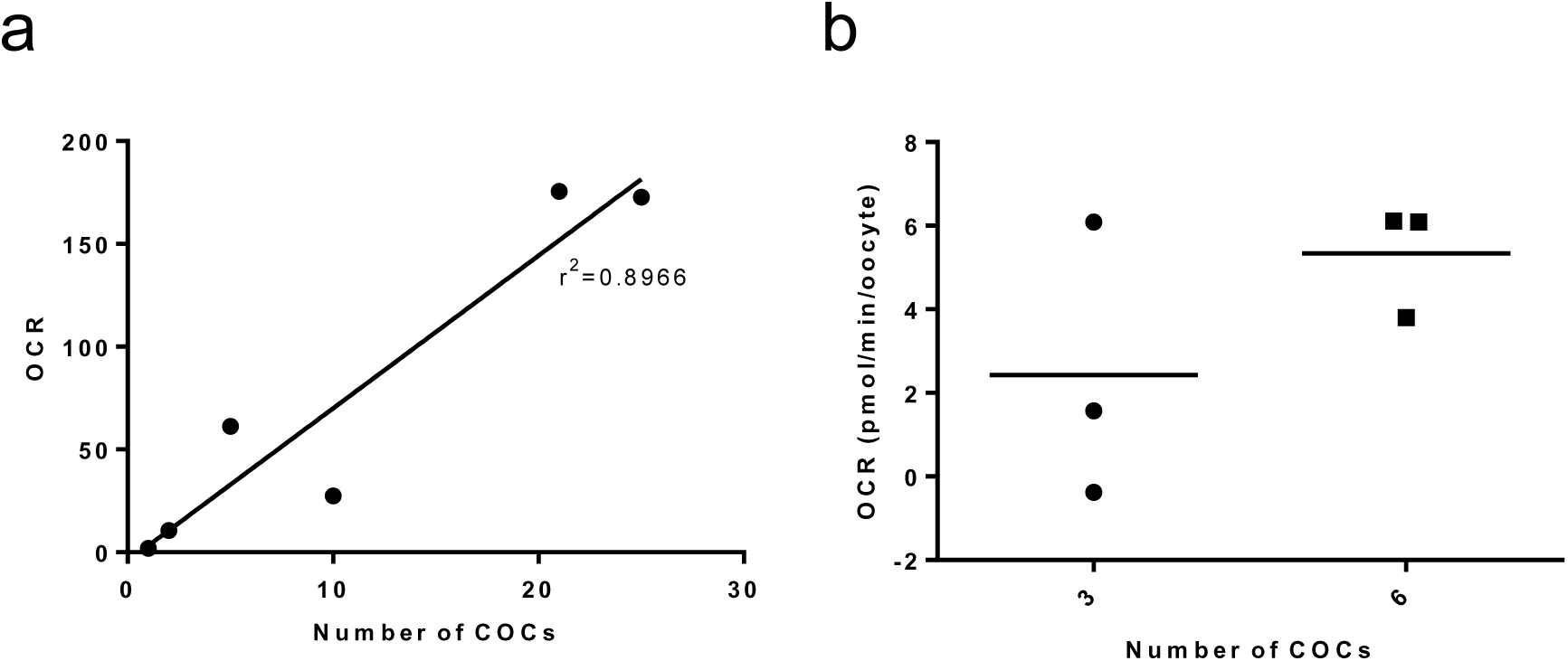
Determination of group size for oxygen consumption measurements in bovine COCs. (a) The linearity of OCR reading for between 1 and 25 oocytes (n=1) which shows significant correlation between number of oocytes per well and measured value of OCR (p=0.0042). (b) A comparison between basal OCR for Seahorse XFp analysis of groups of 3 or 6 COCs (n=1).

**Figure S2.**
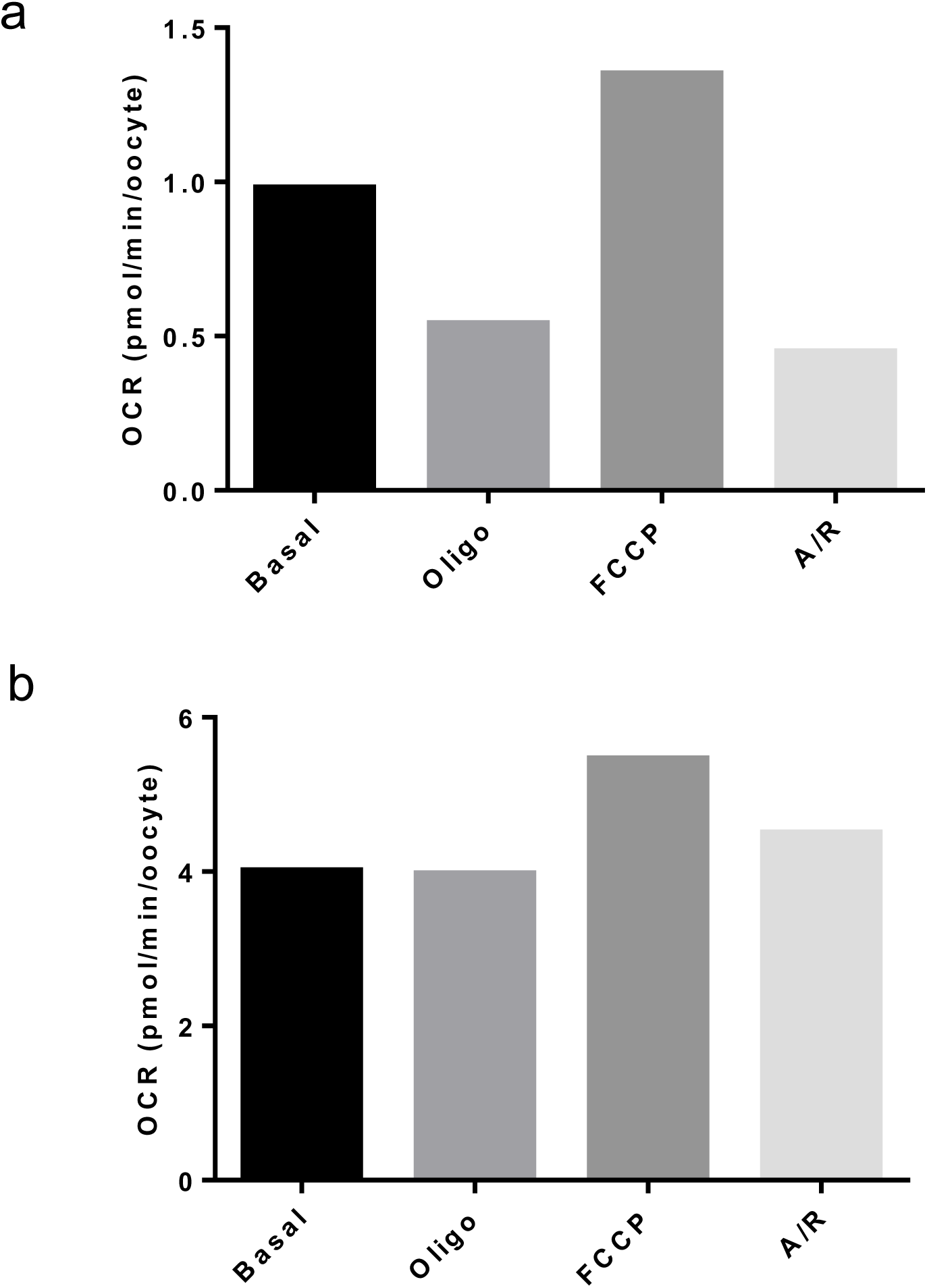
Application of mitochondrial inhibitors to denuded mouse and human M-II oocytes. Mitochondrial inhibitors oligomycin (1 µM), FCCP (5 µM) and antimycin A and rotenone (2.5 µM) were applied in sequence in (a) Mouse oocytes (n=1, representative of 8 oocytes), and (b) Failed-to-fertilise human oocytes (n=1, representative of 5 oocytes). Each data set represents a single data point.

**Figure S3.**
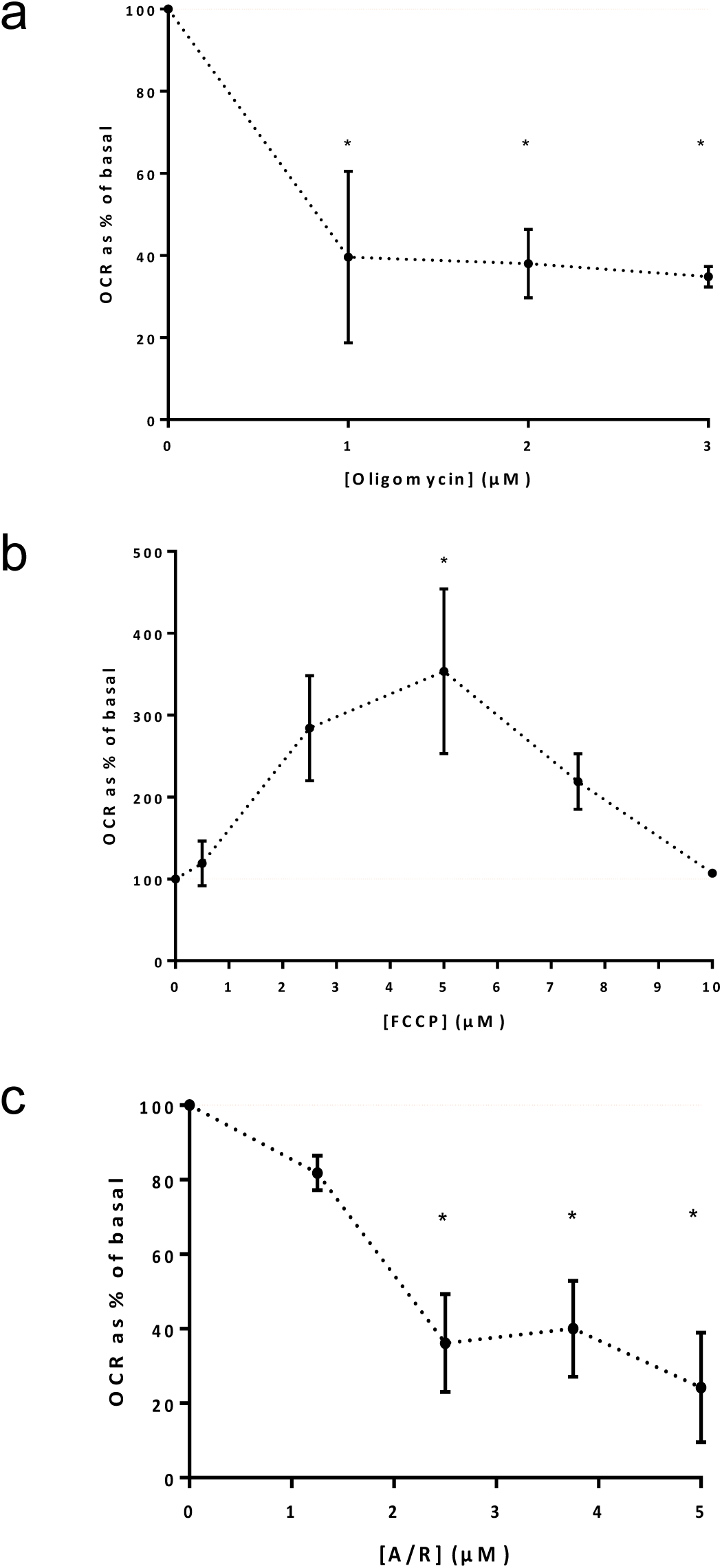
Optimisation of drug concentrations used for dissecting the components of oxygen consumption in bovine oocytes. (a-c) Rate vs. dose plots for inhibitors. Final concentrations chosen were verified in at least two independent experiments. Data shows mean ± SEM. Significance assessed using one-way ANOVA, to p<0.05. Each point is representative of a minimum of two experimental replicates (12 CEOs).

**Figure S4.**
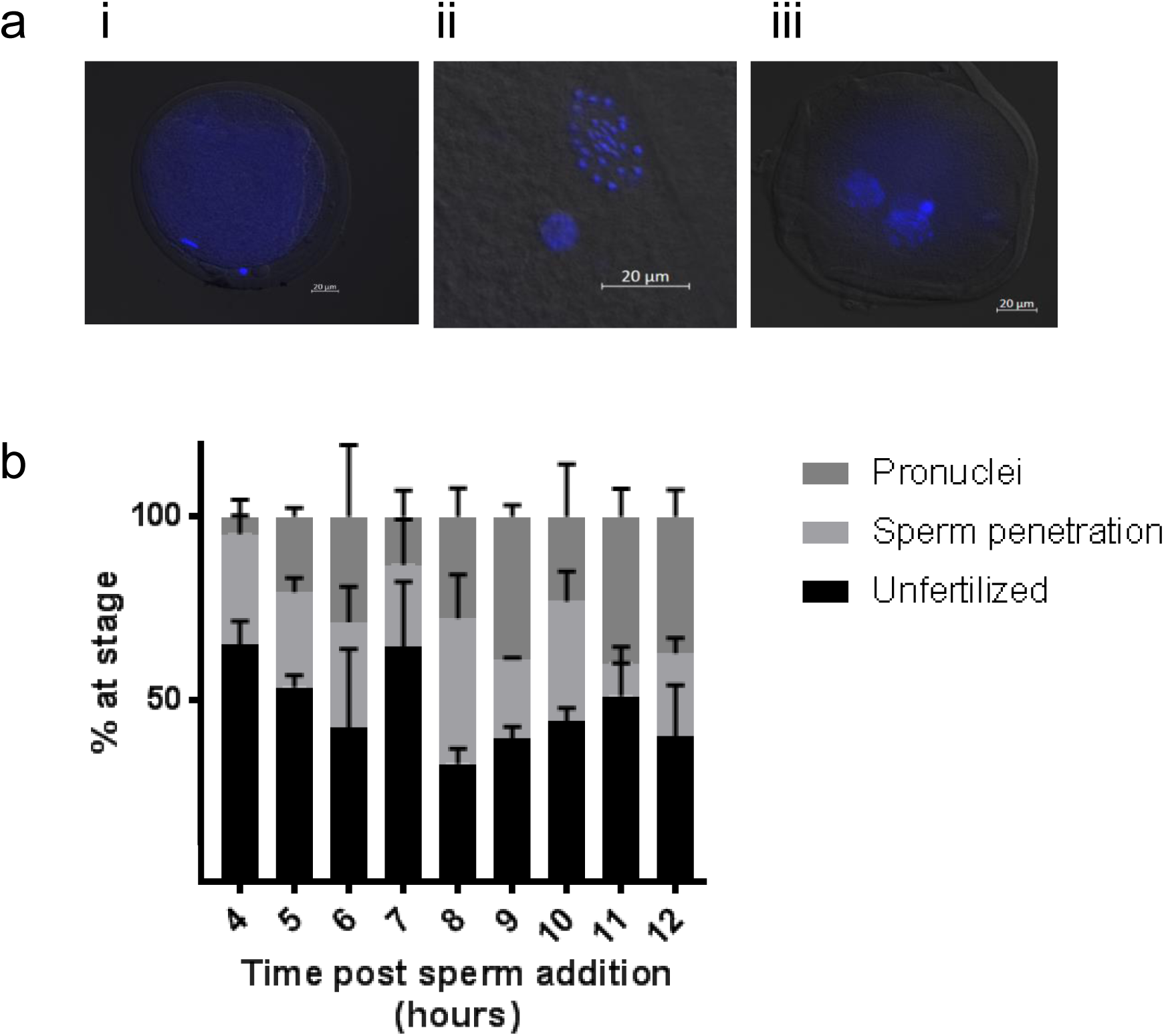
Determination of timing of PN stage in bovine embryos. Hoechst staining was used to determine timing for pronuclear formation in bovine embryos following IVP procedures at 4-12 hours following the addition of motile sperm. (a) Representative images showing Hoechst nuclear staining in bovine zygotes, indicating (i) unfertilized oocyte, (ii) sperm penetration, and (iii) presence of two pronuclei. (b) The breakdown of zygote stage at 4-12 hours post sperm addition. N=3, mean ± SEM, 10 oocytes assessed at each time point per experiment.

**Figure S5.**
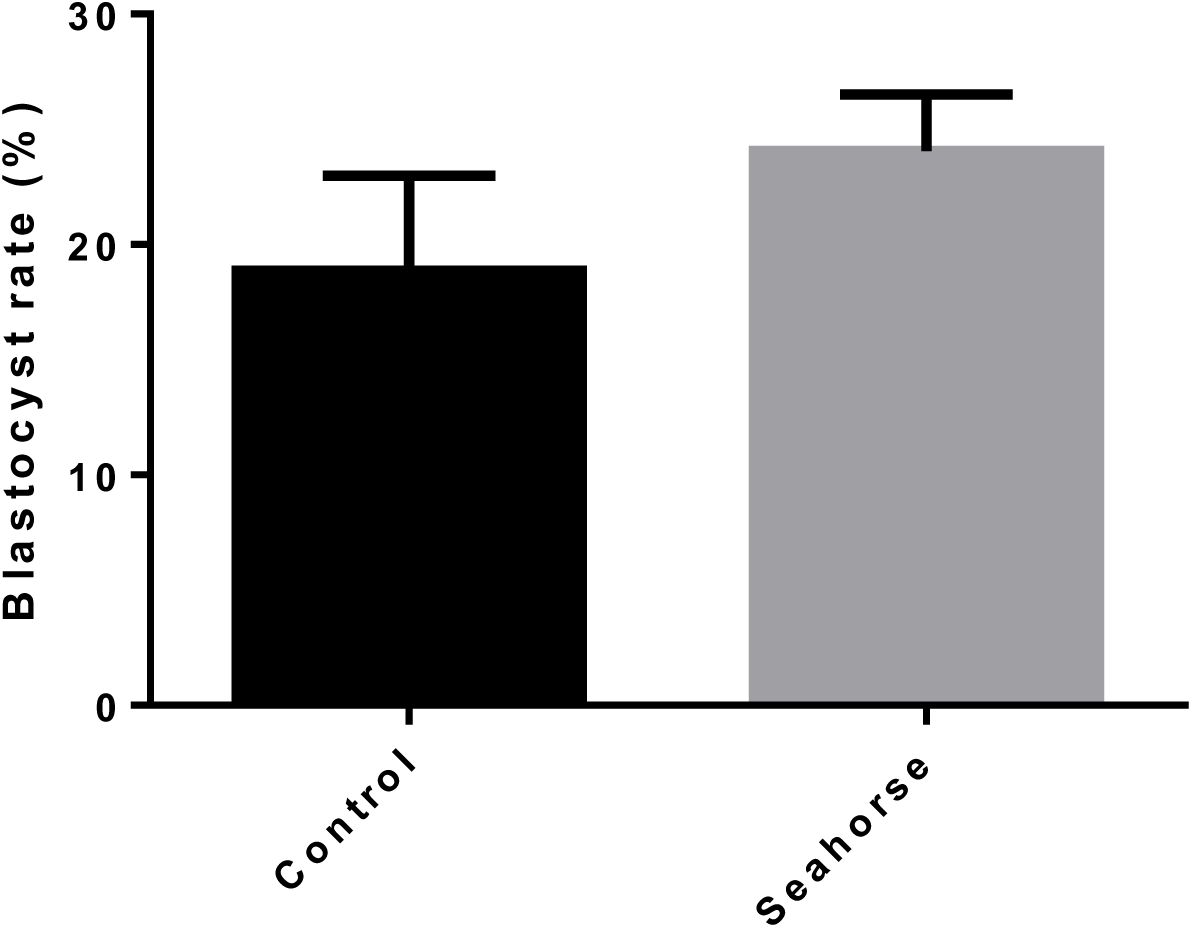
Blastocyst rate following 1 hour EFA on D2. Embryos were either subjected to a 1 hour basal OCR measurement (12 measurements) (Seahorse) or moved into HEPES SOF for the time of the assay (control) before being combined into groups of 15-18 embryos for standard embryo culture. Blastocyst rate was assessed daily from D6 to D8. N=3, representing 6 culture drops per group.

**Figure S6.**
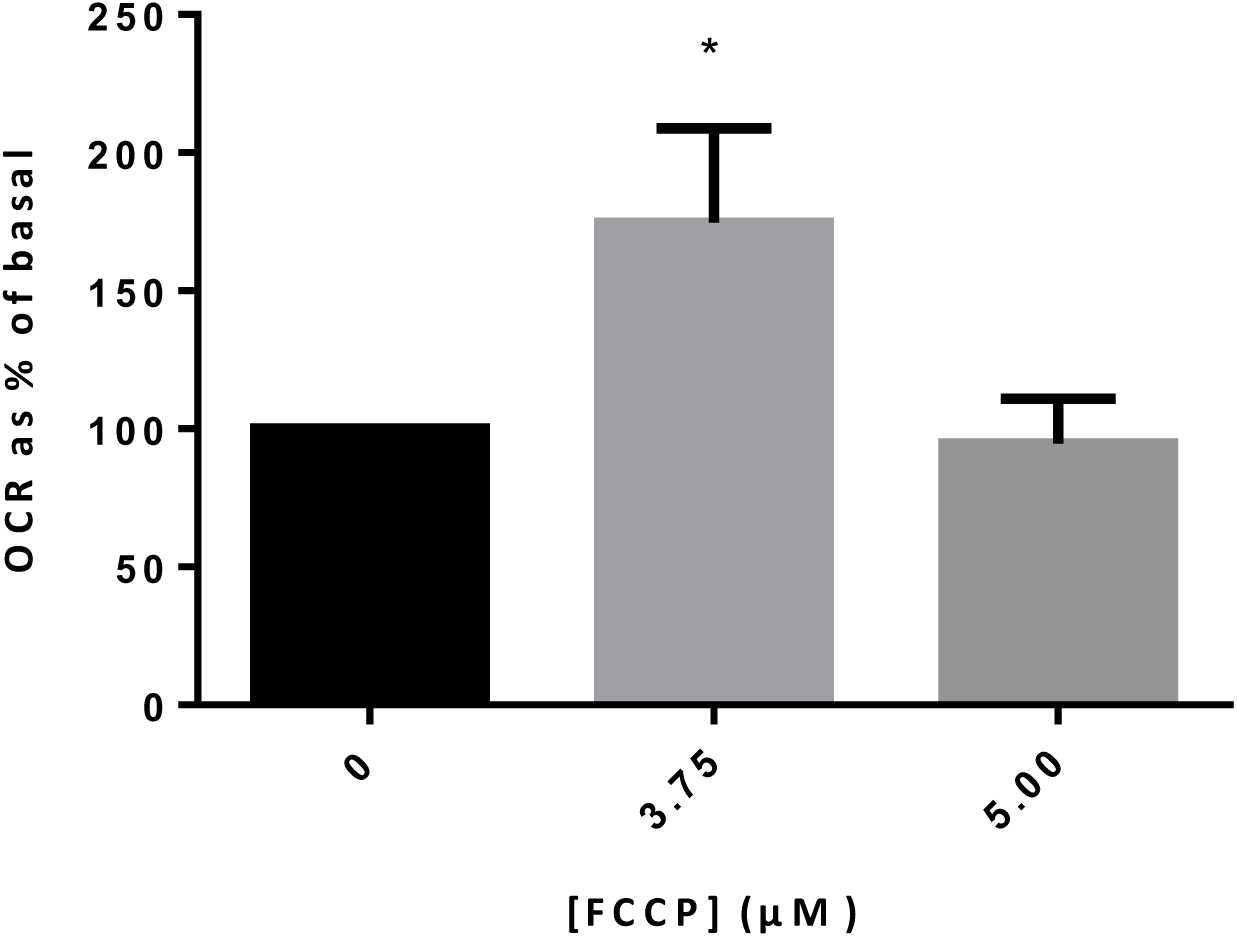
FCCP drug optimization in bovine embryos. FCCP drug optimization at the 2-4 cell cleavage stage. Mean ± SEM (n=3, representative of 18 embryos for each concentration). 100% represents basal, and drug response is shown as a proportion of this. Oligomycin and Antimycin/Rotenone produced the expected responses and thus were used at 1 and 2.5µM respectively. * indicates p<0.05.

**Figure S7.**
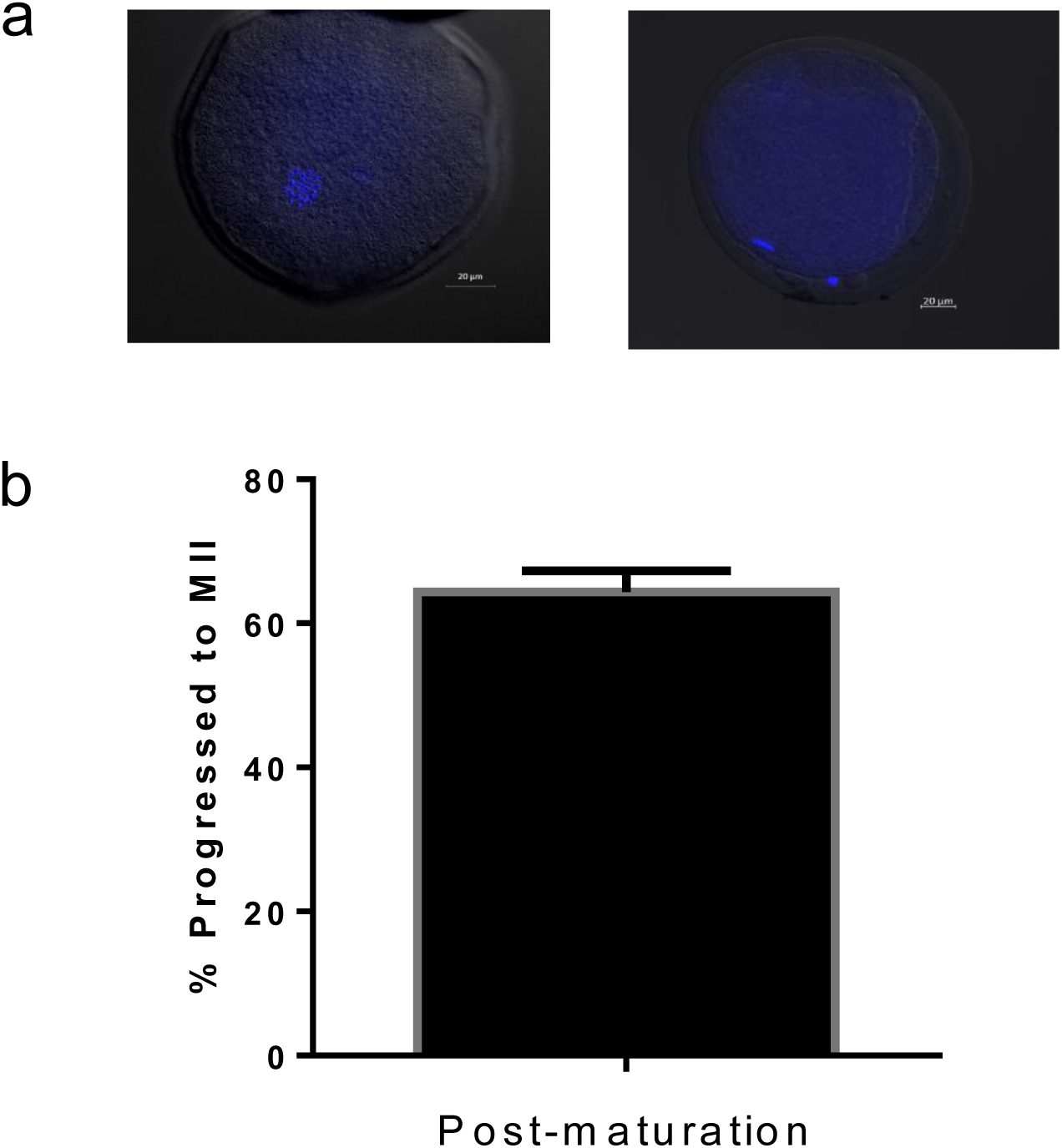
Nuclear status following IVM protocol in bovine COCs. Hoechst staining was applied to determine nuclear status of oocytes following 18-22hr culture in BMM. (a) Representative images showing oocytes at (i) GV and (ii) M-II stage. (b) Percentage of oocytes at M-II stage following IVM. Data represents mean ± SEM (n=3, representing 10 oocytes per experiment).

